# Lipid mediator profile of burn wound healing: Acellular cod fish skin grafts promote the formation of EPA and DHA derived lipid mediators during the first seven days

**DOI:** 10.1101/2021.04.08.438952

**Authors:** Aristotelis Kotronoulas, Sigurður Trausti Karvelsson, Marieke Heijink, Randolph Stone, Martin Giera, Ottar Rolfsson

## Abstract

The use of acellular fish skin grafts (FSG) for the treatment of burn wounds is becoming more common due to its beneficial wound healing properties. In our previous study we demonstarted that FSG is a scaffold biomaterial that is rich in eicosapentaenoic acid (EPA) and docosahexaenoic acid (DHA) conjugated to phosphatidylcholines. Here we investigated whether EPA and DHA derived lipid mediators are influenced during the healing of burn wounds treated with FSG. Deep partial and full thickness burn wounds (DPT and FT respectively) were created on Yorkshire pigs (n=4). DPT were treated with either FSG or fetal bovine dermis while FT were treated either with FSG or cadaver skin initially and followed by a split thickness skin graft. Punch biopsies were collected on days 7, 14, 21, 28 and 60 and analysed in respect of changes to approximately 45 derivatives of EPA, DHA, arachidonic acid (AA), and linoleic acid (LA) employing UPLC-MS/MS methodology. Several EPA and DHA derivatives, including 18-HEPE and 17-HDHA, were significantly higher on day 7 in the DPT when treated with FSG. A similar but non-significant trend was observed for the FT. In addition, prostaglandin F2α and its 15-keto derivative (AA pathway) as well as 13-HODE and 13-HOTrE (LA pathway) were significantly lower in the FSG treated FT. The results suggest that the use of FSG in burn wound treatment can alter the formation of EPA and DHA mono hydroxylated lipid mediators in comparison to other grafts of mammalian origin. The differences observed during the first seven days after treatment indicates that FSG affects the early stages of wound healing.

**Highlights:**

- This is the first study to investigate whether omega-3 rich biomaterial such as fish skin graft can affect the lipidome of burn wound healing
- The study monitors the formation of lipid mediators during 60 days of burn wound healing
- This is the first study to show an increase in the formation of mono-hydroxylated derivatives of EPA and DHA after seven days of treatment with fish skin graft
- The study showed significantly decreased formation of lipid mediators involved in pain modulation in wounds treated with fish skin graft

## 1. Introduction

Skin grafting is an important part of the therapy of burn wounds and its early application can avoid complications including sepsis, multi-organ failure, and acute kidney injury.[1] Excision of the burned area and grafting with autologous split thickness grafts (STSG) currently represents the standard of care of deep partial and full thickness burn wounds. Lack of availability and scar formation are however disadvantages associated with the use of STSG at present and have encouraged the search and development of the ideal skin substitute.[2] These materials include allografts and xenografts from different animal origins.[3] Currently, acellular cod fish skin grafts (FSG) from the North Atlantic cod (Gadus morhua) are increasingly used in hospitals with promise of being an effective alternative tissue source for the wound care of a variety of etiologies.[4] FSG has been successfully applied to the treatment of acute [5, 6] and hard to heal wounds,[4] providing a readily safe skin substitute, free of the risk of transmission of viral disease and auto-immune reaction risk due to its piscine, cold water origin.[4, 7] Recent studies have shown that the FSG benefits wound healing on account of its ability to support cell ingrowth of both fibroblasts and adipose stem cells [8] with concomitant pain relief and reduced scarring.[6, 7]

The four principal phases of the wound healing process (e.g., hemostasis, inflammation, proliferation, and remodeling) are orchestrated by a complex series of biochemical changes where different compounds classes (e.g., amino acids, lipids, cytokines, growth factors, endocannabinoids, and others) play an important role.[9, 10] The inflammatory phase and its resolution are considered crucial for the favorable advancement to wound healing.[10, 11] Several pro-inflammatory lipid mediator derivatives of arachidonic acid (AA) and linoleic acid (LA), such as the groups of prostaglandins, leukotrienes, hydroxy-, and keto-eicosatetraenoic acids (HETEs and KETEs respectively) are important in this regard.[11] In addition, omega-3 fatty acids (ω3-FA) such as eicosapentaenoic acid (EPA) and docosahexaenoic acid (DHA) can promote wound healing by resolving inflammation through the formation of specialized pro-resolving lipid mediators (SPM).[12-15] During inflammation, the different types of SPMs (resolvins, protectins, and maresins) are biosynthesized in exudates by immune cells and/or via transcellular biosynthesis involving endothelial or epithelial cells.[14] They control the resolution of inflammation by halting excessive infiltration of neutrophils and stimulating the expression of genes important for antimicrobial defense,[14, 16] diminishing the production of pro-inflammatory lipid mediators and chemokines [17] and promoting wound re-epithelialization.[18]

In a recent study, our group demonstrated that FSG is particularly rich in phosphatidylcholines that contain EPA or DHA and that this differentiates FSG from other skin grafting materials.[19] Thus, it has been postulated that the use of ω3-FA enriched scaffold biomaterials can be an effective way to topically deliver ω3-FA to wounds and enhance the healing processes via resolution of inflammation. The goal of the present study was to explore whether the FSG derived ω3-FA can alter the formation of lipid mediators during burn wound healing, as opposed to other grafts of mammalian origin that are not rich in ω3-FA. For this, deep partial and full thickness burn wounds (DPT and FT respectively) were created on Yorkshire pigs and treated with different grafts treatments. Wound biopsies were then collected during the first 60 days of healing. Clinical and histological data and evaluation of the wound healing process of the DPTs have already been published elsewhere.[20] In this study, the changes in lipid mediators were monitored by liquid chromatography–mass spectrometry (LC/MS) during the healing period and their rate of formation was compared between wounds treated with the different grafting materials.

## 2. Materials and Methods

### 2.1 Chemical and reagents

Synthetic standards of lipid mediators were purchased from Cayman Chemical (Ann Abor, Michigan, USA). Water used for sample preparation was obtained from Biosolve (Valkenswaard, The Netherlands), mobile phase water was from Honeywell Riedel-de Haën (Seelze, Germany). LC-MS grade MeOH was obtained from Merck (Darmstadt, Germany). Formic acid and *n*-hexane were obtained from VWR (Darmstadt, Germany). Acetic acid was purchased from Fluka (Darmstadt, Germany).

### 2.2 In vivo experiment

All experiments were carried out under a US Army Institute for Surgical Research (USAISR) approved IACUC protocol (A-16-021-TS5 approved May 5, 2017). Research was conducted in compliance with the Animal Welfare Act, the implementing Animal Welfare Regulations, and the principles of the Guide for the Care and Use of Laboratory Animals, National Research Council. The facility’s Institutional Animal Care and Use Committee approved all research conducted in this study. The facility where this research was conducted is fully accredited by AAALAC. Animals were housed individually in a temperature-controlled environment with a 12-h light/dark cycle in the AAALAC approved vivarium at the USAISR with access to water and food ad libitum. A schematic overview of the *in vivo* experiment can be found in **Figure 1**. Before burning, hair was removed from the dorsum of the animals by chemical depilation (Nair(tm)) was rinsed with sterile water. Four DPT and FT (5×5 cm) were created on the dorsum of anesthetized Yorkshire pigs (n=4) using appropriate pain control methods. The burn depth and treatment on the back of the pig were randomized. Burns were then excised down to a bleeding wound bed twenty four hours post-burn. DPT were then treated with either omega-3 rich FSG (treatment 1, n=2 for each pig) or fetal bovine dermis graft (treatment 2, n=2 for each pig). A reapplication of the FSG was applied after 7 days, and all wounds were allowed to heal by secondary intentions. Fetal bovine dermis xenografts used were commercially available under the trademark Primatrix (TEI Biosciences Inc, Boston, Massachusetts). On the other hand, FT were first treated with either FSG (treatment 3, n=2 for each pig) or cadaver skin (treatment 4, n=2 for each pig) until day 7 when the wounds from both treatment 3 and 4 were grafted with a 1.5:1 meshed split thickness skin graft.

**Figure 1:**
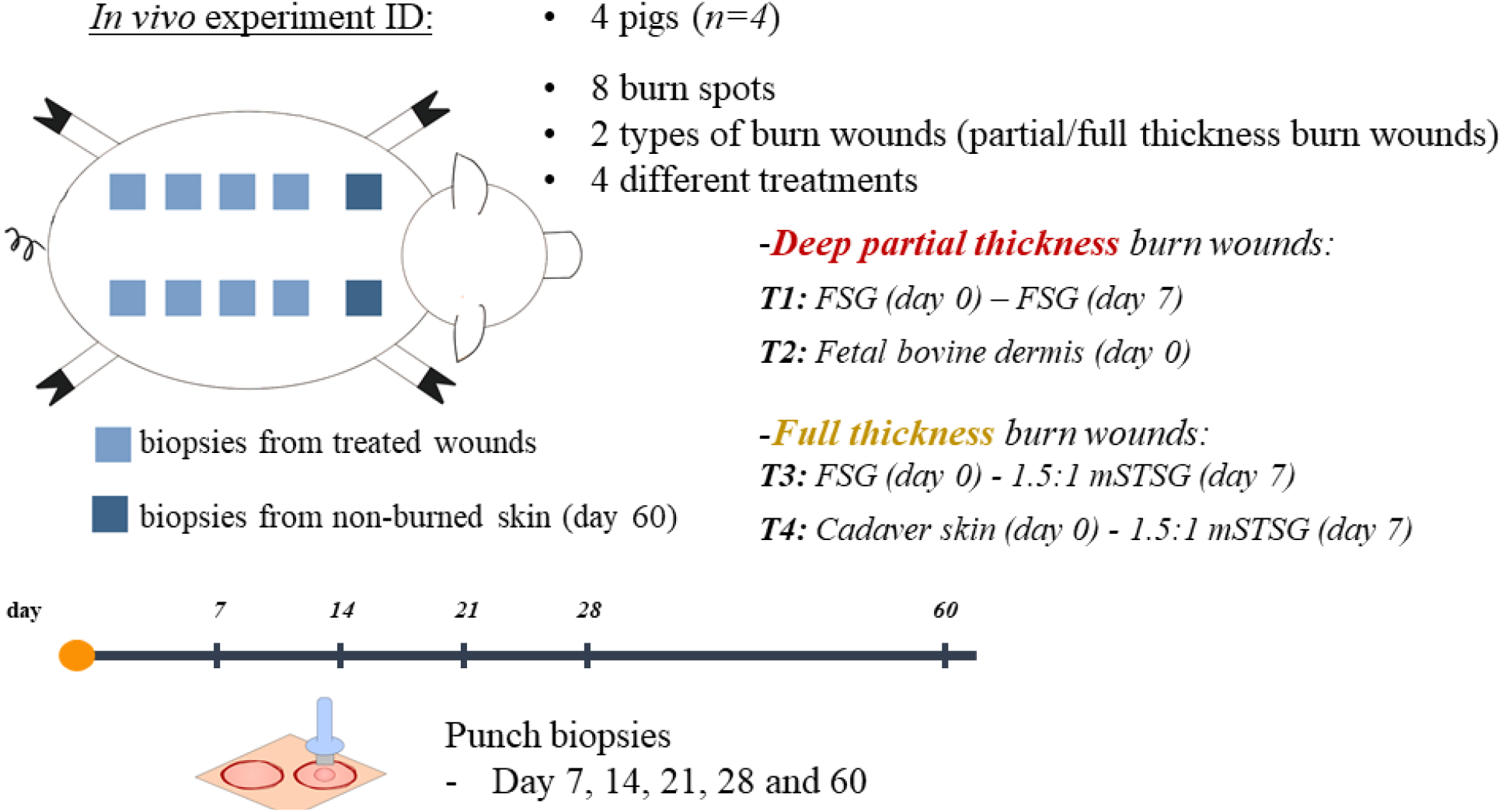
Experimental procedure and sampling. 5×5 cm partial (n=4) or full thickness (n=4) burn wounds were created on the dorsum of anesthetized Yorkshire pigs (n=4). The wounds received different treatment procedures (T1-4) based upon their severity. T1 or T2 were applied to the deep partial thickness wounds and T3 or T4 were applied to the full thickness wounds. For T1, FSG graft was applied immediately after the cleaning and debridement of the wound and a second FSG graft was applied seven days after. For T2, a fetal bovine dermis graft was applied to the wound immediately after the cleaning and debridement of the wound and was left there until the wounds were healed. For T3, an FSG graft was applied followed by a meshed split thickness skin graft (mSTSG) on D7. For T4, cadaver skin graft was applied followed by the application of mSTSG on day seven. Punch biopsies were collected on days 7, 14 21, 28, and 60 after the application of the corresponding graft. On day 60, two punch biopsies were also collected from a non-burned part of the pig’s dorsum

Cadaver skin was acquired by harvesting skin grafts with a pneumatic dermatome from recently euthanized Yorkshire pigs. The areas that were used to acquire the Cadaver skin were on the back, sides, and hindlimb areas. Allografts were soaked in sterile saline with penicillin/streptomycin (pen/step) for ∼1 hr, 0.1% benzalkonium bromide for ∼15 min, washed 3 times in normal saline with pen/strep, and were then stored in 85% glycerol at 4°C until the day of the experiment. On the day of experiment, the allografts were rinsed with normal saline for up to 1 hr to remove any preservation residue and return to room temperature which allowed easier application to the wound bed (PMID: 16023938 and 16023931).

Autologous skin was harvested from the hindlimbs with the dermatome set to 0.012”. Autografts were meshed to 1.5:1 to create the meshed split thickness skin graft (PMID: 27600980 and 29457376). Rechecks were performed on days 7, 14, 21, 28, 45, and 60 during which digital images, non-invasive measurements, and punch biopsies were acquired and immediately frozen at -80°C. In addition, on day 60, biopsies were collected from two spots other than the wounds (non-burned skin, nb60) in order to be used as control for the healthy skin.

### 2.3 Sample extraction

The punch biopsies were thawed at room temperature, weighed, and subsequently homogenized using plastic pestles in 0.5 mL of 70 % MeOH for 1 min. Following centrifugation at 13000 rpm for 10 min the supernatant was set aside. The homogenization process was repeated on the pellet with another 0.5 mL of 70% MeOH and the supernatants were combined and immediately frozen at -80°C until further analysis.

### 2.4 UPLC-MS/MS Analysis

#### 2.4.1 Sample preparation and purification

The combined supernatants were purified by solid phase extraction. To the supernatants 4 µL of internal standard mix (containing 8-iso-PGF_2α_-d4, PGE_2_-d4, LTB_4_-d4, 15-HETE-d8 and DHA-d5) was added. Subsequently, the supernatant was diluted with water, acidified using formic acid and applied to Sep-Pak C18 cartridges (200 mg, 3cc, Waters, Milford, MA, USA) that had been conditioned with MeOH and equilibrated with water. Following washing with water and *n*-hexane the samples were eluted using methyl formate. The eluate was dried under a gentle stream of nitrogen and reconstituted in 40% MeOH.

#### 2.4.2 Instrument specifications

The LC system consisted of a Shimadzu SIL-30AC autosampler, two LC-30AD pumps and a CTO-20AC column oven. Forty µL sample was injected. Separation was achieved on a kinetex C18 column (Phenomenex, Aschaffenburg, Germany, 50 × 2.1 mm, 1.7 µm) with a gradient of 0.01% acetic acid in water (solvent A) and 0.01% acetic acid in MeOH (solvent B). The flow rate was 400 µL/min. The gradient was as follows: 0.0-1.0 min. isocratic at 30% B, 1.0-1.1 min. linear increase to 45% B, 1.1-2.0 min. linear increase to 53.5% B, 2.0-4.0 min. linear increase to 55.5% B, 4.0 – 7.0 min. linear increase to 90% B, 7.0 – 7.1 min. linear increase to 100% B, 7.0-9.0 min. isocratic at 100% B, 9.0–9.5 min. linear decrease to 30% B, 9.5-11.5 min. isocratic at 30% B. Analytes were detected on a Sciex Qtrap 6500 equipped with an ESI-source and operated in negative scheduled MRM mode. The settings of the source were as follows: needle voltage -4500V, drying temperature 450 °C, gas 1/nebulizer gas (air) 40 psi, gas 2/drying gas (air) 30 psi, nebulizer gas (nitrogen) 30 psi. The entrance potential was set to -10V and the collision gas to ‘medium’. The individually optimized parameters for the individual compounds are summarized in the supporting information. A summary of the detected analytes included is shown in Table 1. The compounds m/z transitions used for their detection are summarized in the supporting information.

**Table 1:**
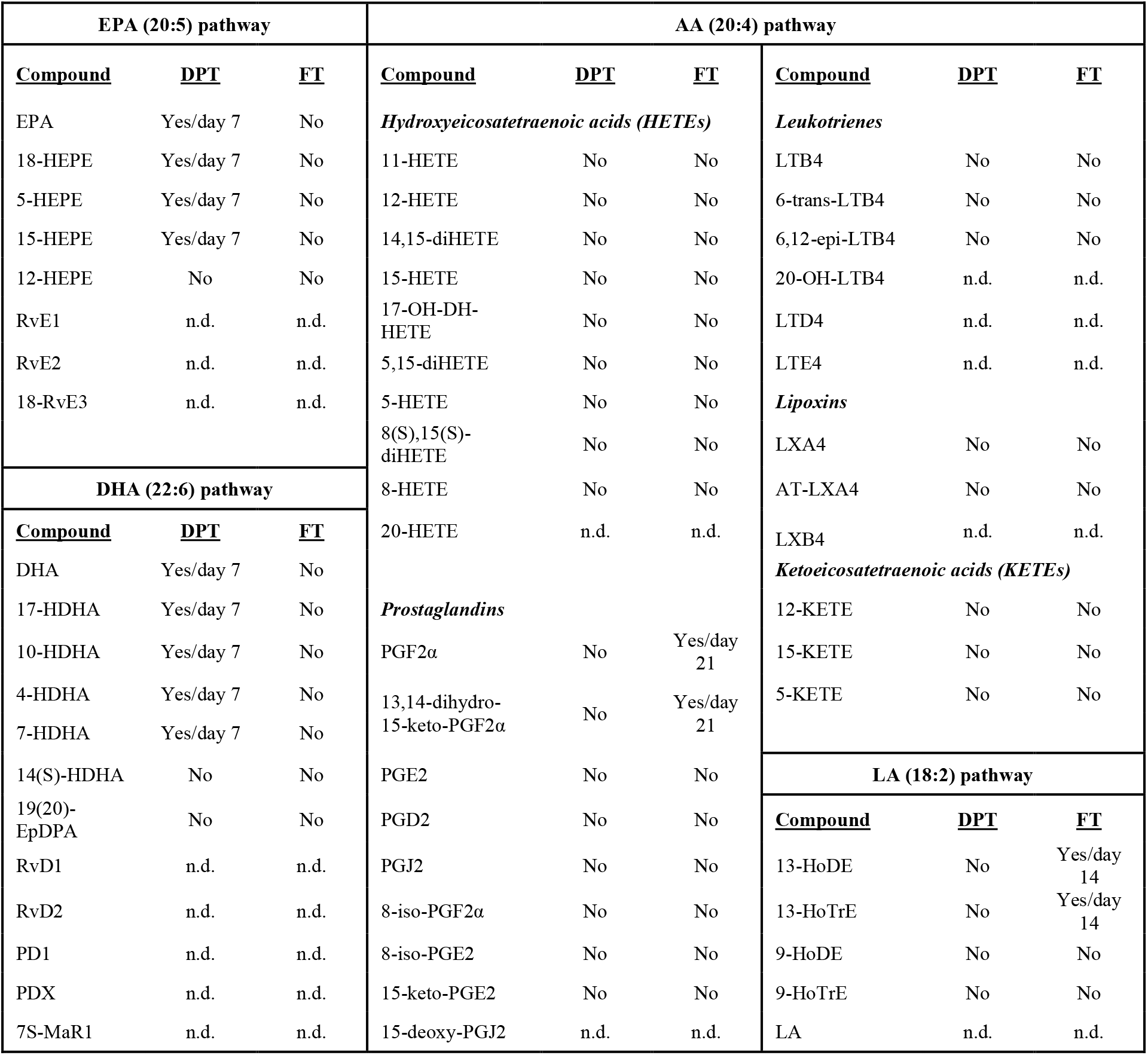
Statistical analysis of lipid content following deep partial and full thickness burns. Summary of all lipid mediators measured with UPLC-MS/MS. The method included 60 compounds from which 15 were not detected (n.d.) in the samples. For the detected lipid mediators, the average normalized signal intensities were calculated at each time point in each type of wound and compared between treatments. (Yes): significant difference (p<0.05) was observed between treatments. The time point that the difference was observed is annotated as day 7, 14 or 21. (No): no significant differences between treatments was observed at any time point.

### 2.6 Normalized signal intensities and statistical analysis

For the detected compounds, the measured signal intensities were normalized based on the corresponding internal standards and weight of biopsy in order to acquire the normalized signal intensities (NSI). After that, the average NSI was calculated for each compound and plotted for the different time points, type of wounds, and received treatment. The generated plots for all detected metabolites can be found in supplementary information. The average NSI between groups at each time point were compared using student t-test and was carried out with IBM SPSS Statistics for Windows, Version 24.0. Armonk, NY: IBM Corp.

## 3. Results

### 3.1 EPA and DHA are increased in FSG treated wounds

In order to explore the formation of lipid mediators during the first 60 days of wound healing and detect possible changes in their concentrations, 60 derivatives of EPA, DHA, AA, and LA and free fatty acids that are part of their metabolic pathways, were analyzed by UPLC-MS/MS. A summary of the analytes included in the method is shown in **Table 1**. As shown, 45 different lipid mediators were detected in all biopsies, independently of the treatment.

Focusing on these compounds, the acquired average normalized signal intensity (NSI) was calculated for the different time points, type of wounds, and received treatment. General trends were visualized using a heatmap displaying fold changes as compared to non-burned skin collected in day 60 (**Figure 2**). In the DPTs treated with FSG (Treatment 1) (**Figure 2A**), the EPA and DHA derived mediators increased as early as day 7, as opposed to the fetal bovine dermis xenografts (treatment 2) treated wounds (**Figure 2B**) where the derivatives of these compounds increased at later stages during healing or around day 14 and 21. As opposed to EPA and DHA derivatives, lipids of the AA and LA pathways in the DPTs showed a later increase (day 14 and 28), and decreased around day 28 and 60, independently of the treatment.

**Figure 2:**
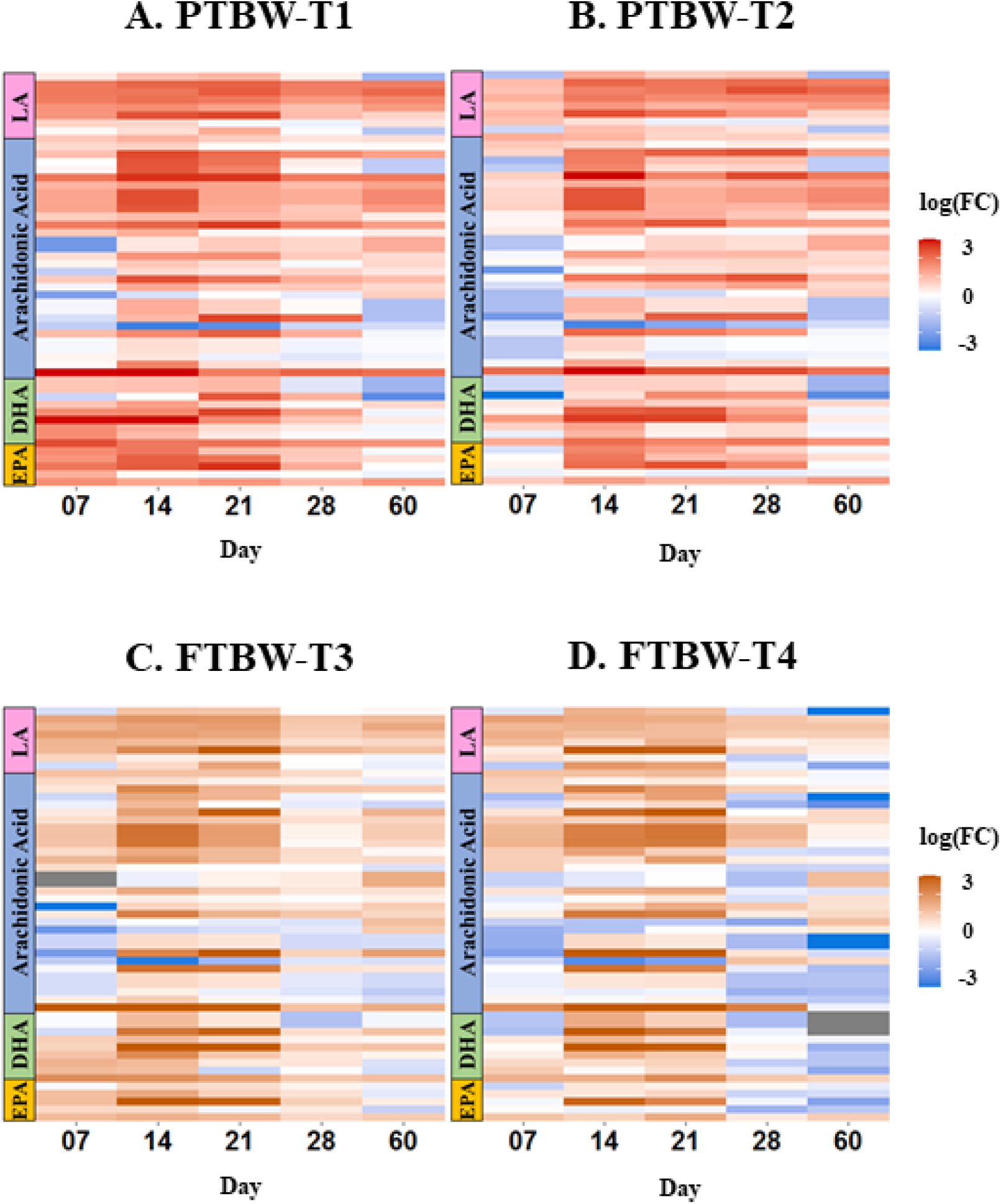
Changes to lipid content are observed during burn wound healing irrespective of treatment. Changes to the average signal intensity of each monitored lipid at the different sampling time points with respect to non-burned skin at day 60. The change in average normalized signal intensity for all detected lipid mediators over time is presented in heatmaps for each individual treatment. A summary of the lipids in the experimental method, whether they were detected or not and significant differences (p<0.05) observed between samples of different treatment for the same time point are reported in **Table 1**. Grey color indicates that the compound was not detected on the specific day.

A similar behavior was observed for the FT (**Figure 2C**), where the EPA and DHA derivatives in the FSG (treatment 3) treated wounds had increased on day 7 as opposed to the biopsies from the cadaver skin treated wounds (**Figure 2D**) where their values appeared higher from day 14 and 28. Derivatives of AA and LA increased at days 14 and 21 independent of treatment. Finally, most of the detected compounds in FT decreased at days 28 and 60, independent of treatment. These analyses show that FSG modulates the lipid profiles of burn wound healing in both DPTs and FTs.

The average NSI were plotted individually for all detected metabolites at the different time points with respect to the type of wound (**Figure 3**). EPA and DHA **Figure 3**showed an increasing trend in the FSG treated wounds (treatment 1 or treatment 3) as opposed to the other treatments and this trend was independent of the wound type. However, the statistical analysis showed that these changes were only significant (*p<0*.*05*) at day 7 for DPTs **(Figure 3A)**. The average NSI of EPA on day 7 was 0.063±0.032 vs 0.027±0.028, for treatment 1vs 2, respectively (*p=0*.*03*) and of DHA was 0.085±0.038 vs 0.05±0.004, for treatment 1 vs 2, respectively (*p=0*.*0004*). In the following days, the average NSI values showed no significant differences between treatments.

**Figure 3:**
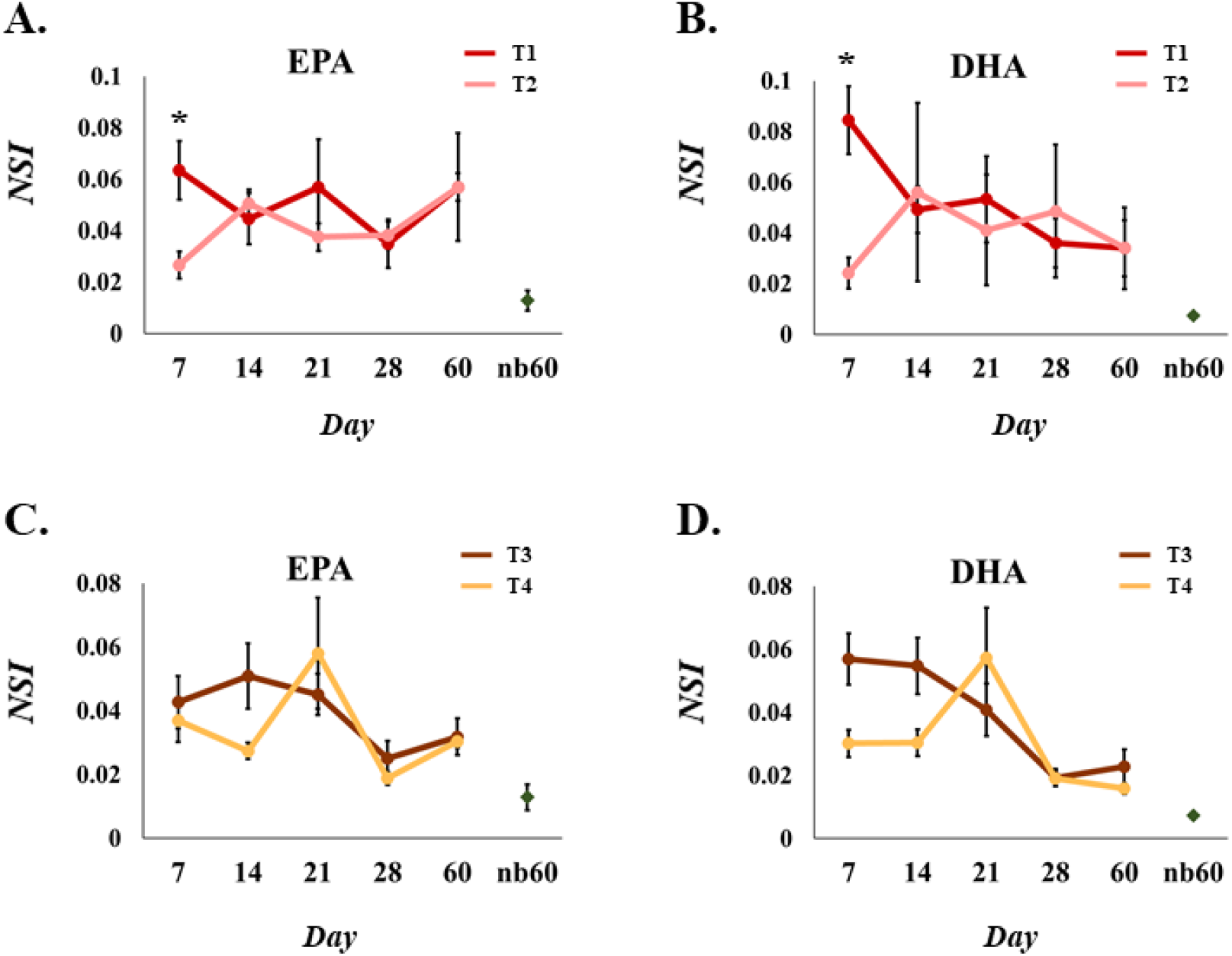
EPA and DHA profiles change during burn wound healing. EPA and DHA derived lipid mediator profiles acquired overs 60 days of wound healing for deep partial thickness burn wounds (A and B) and full thickness burn wounds (C and D). Intensities of lipids are displayed as average normalized signal intensities (NSI). Nb60 represents the values determined in the non-burned skin samples harvested at day 60. *Significant differences (p<0.05) between treatments in the corresponding time point. The error bars show the standard error of the mean.

As opposed to the DPTs, no statistically significant differences between treatments at any of the time points for either EPA or DHA were detected in the full thickness burn wounds (**Figure 3B**). DHA was however on average higher in the fish skin treatment 3 treated FTs than cadaver skin treated burns at day 7 (0.057±0.020 in treatment 3 and 0.030±0.014 in treatment 4) and at day 14 (0.055±0.022 in treatment 3 and 0.030±0.010 in treatment 4). These differences however were non-significant between treatments due to high standard deviations. Nevertheless, a trend towards higher amounts of both EPA and DHA was observed in the treatment 3 treated wounds. In summary, the topical treatment of burn wounds with fish skin results in changes to EPA and DHA lipid profiles during burn wound healing. Next, we focused on the changes to EPA and DHA derived lipid mediators.

### 3.2 Derivatives of EPA and DHA are increased in FSG treated wounds DPTs

A similar type of analysis was followed to investigate changes to lipid mediator derivatives of EPA and DHA. The calculated average NSI for the measured lipid mediators were plotted for the different time points for both DPT and FT. A summary of the observed differences for each of the monitored compound are reported in **Table 1**.

For the DPT, most of the detected EPA and DHA lipid mediators showed significant differences at day 7. As shown in **Figure 4 A-G**, the average NSI values were significantly higher in FSG when compared to fetal bovine dermis xenografts for 18-HEPE (*0*.*063±0*.*040 vs 0*.*015±0*.*012*, treatment 1 vs 2 *respectively, p=0*.*007*), 5-HEPE (*0*.*028±0*.*025 vs 0*.*006±0*.*005, p=0*.*03*), and 15-HEPE (*0*.*026±0*.*025 vs 0*.*004±0*.*02, p=0*.*03*) of the EPA pathway as well as 17-HDHA (*0*.*015±0*.*014 vs 0*.*003±0*.*002*, treatment 1 vs 2 *respectively, p=0*.*04*), 4-HDHA (*0*.*065±0*.*034 vs 0*.*027±0*.*017, p=0*.*02*), 7-HDHA (*0*.*058±0*.*047 vs 0*.*008±0*.*008, p=0*.*01*) and 10-HDHA (*0*.*066±0*.*043 vs 0*.*011±0*.*004, p=0*.*03*) of the DHA pathway. Following day 7, these compounds followed a similar trend, with no significant differences observed between treatments. No other EPA/DHA derived lipid mediators showed significant differences at any of the time points in the DPT.

**Figure 4:**
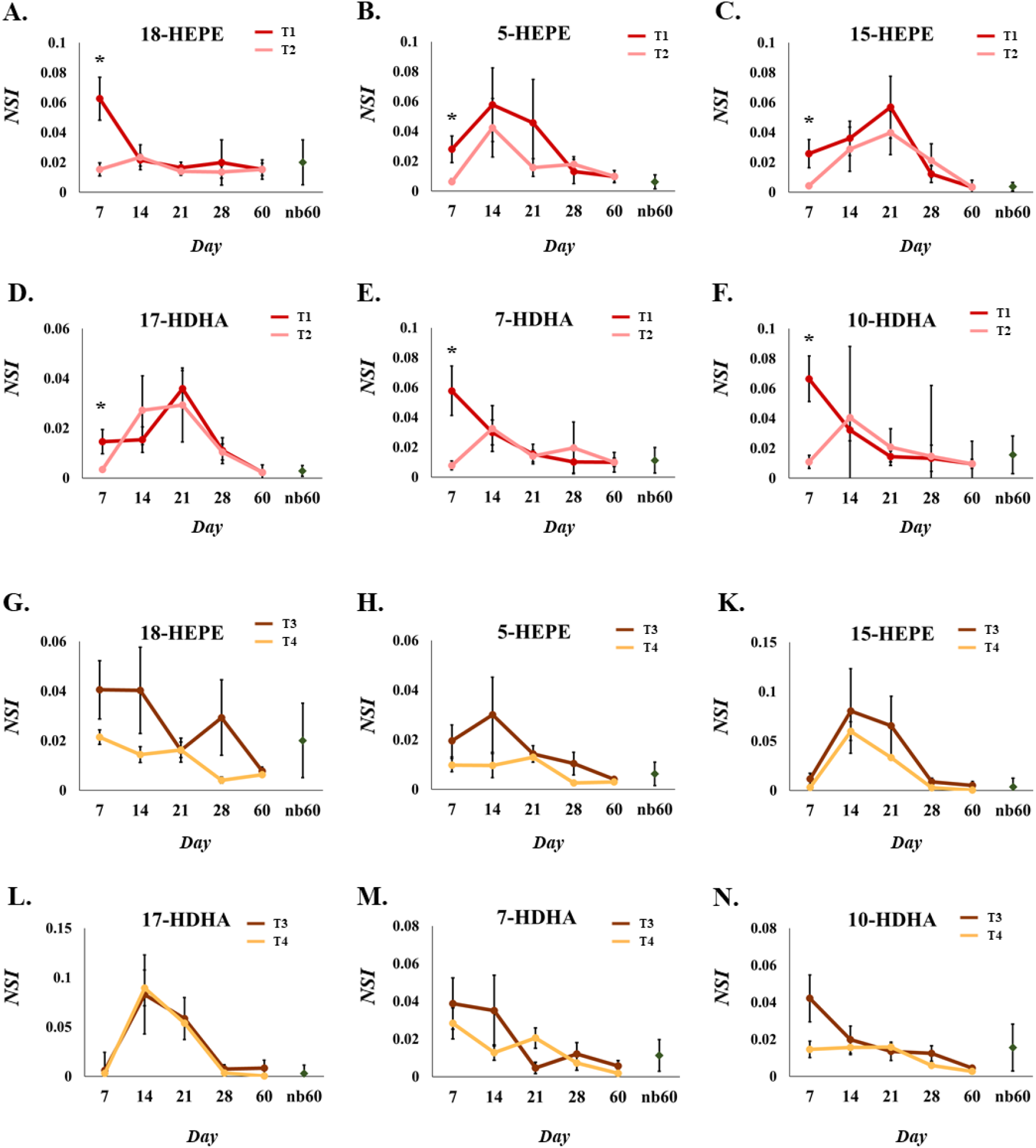
The EPA and DHA derived lipid mediators 18-HEPE and 7-HDHA change during burn wound healing. EPA and DHA derived lipid mediator profiles acquired overs 60 days of wound healing for deep partial thickness burn wounds (A-F) and full partial thickness burn wounds (G-N). Intensities of lipids are displayed as average normalized signal intensities (NSI). Nb60 represents the values determined in the non-burned skin samples harvested at day 60. *Significant differences (p<0.05) between treatments in the corresponding time point. The error bars show the standard error of the mean.

In the case of the FT biopsies, no significant differences between treatments at any of the time points were detected in EPA and DHA derivatives (**Table1**). Trends towards increased EPA and DHA lipid mediator derivatives however were observed in the FSG treated wounds as opposed to the cadaver skin (**Figure5-M**) but due to high standard deviations these were not significant.

For both DPT and FT, the EPA and DHA derived lipid mediators generated plots not shown in **Figure 4** can be found in the supplementary information.

### 3.3 Derivatives of AA and LA are increased in FSG treated wounds FTs

Further analysis of the measured lipid mediators derived from AA and LA did not present any significant differences in the DPT independent of the treatment (**Table1**). In the FT wounds however, PGF2α and 13,14-dihydro-15-keto-PGF2α from the AA pathway and 13-HoDE and 13-HoTrE from the LA pathway were significantly different following treatment at day 21 or day 14 respectively for the AA and LA derivatives (**Table 1 and Figure 5**). As shown, the average NSI on day 21 of PGF2α (*0*.*019±0*.*015 vs 0*.*085±0*.*068, p=0*.*04*) and its keto derivative 13,14-dihydro-15-keto-PGF2α (*0*.*013±0*.*010 vs 0*.*040±0*.*024, p=0*.*03*) were significantly lower for the wounds treated with treatment 3 as opposed to those treated with treatment 4. Similarly, the average NSI on day 14 for 13-HoDE (*0*.*043±0*.*034 vs 0*.*136±0*.*049, p=0*.*006*) and 13-HoTrE (*0*.*044±0*.*038 vs 0*.*135±0*.*068, p=0*.*03*) were significantly lower in the FT treated with treatment 3. The average NSI values for the other time points presented no significant differences between treatments. The generated plots for all lipid mediators that are not shown in **Figure 3, 4, or 5** can be found in the supplementary information.

**Figure 5:**
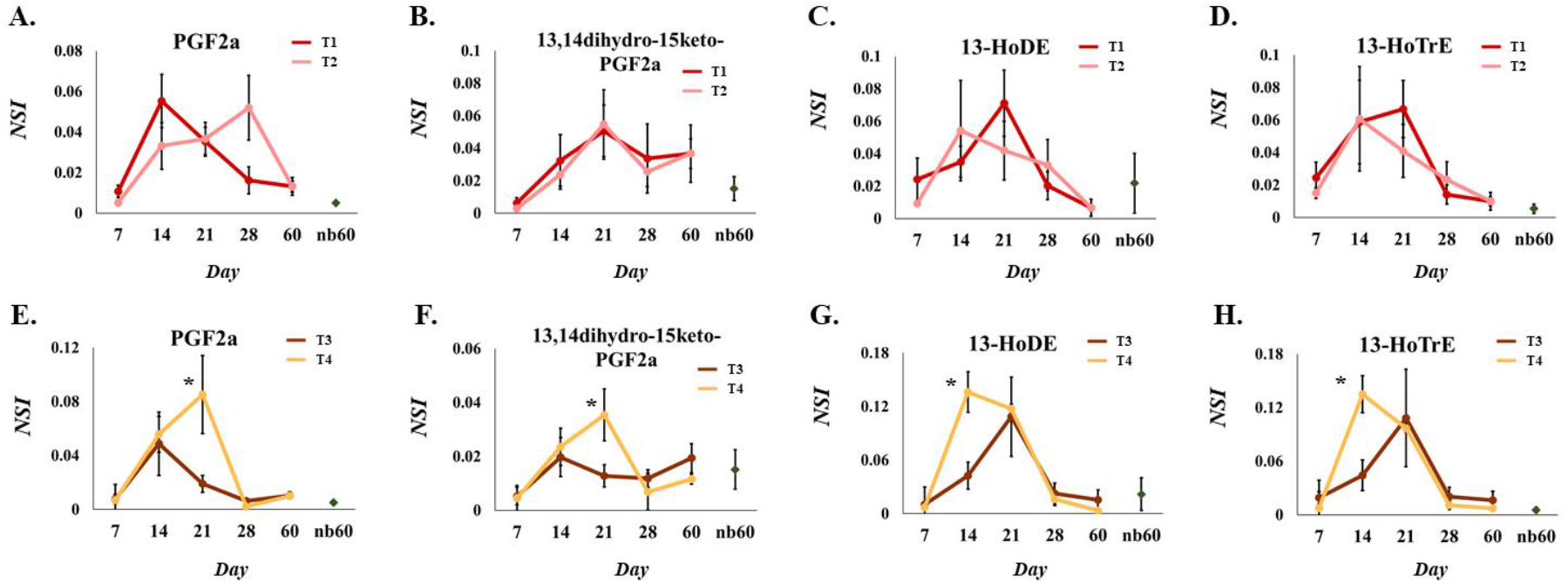
PGF2α and 13-HoDe concentration changes in full thickness burn wounds treated with fish skin. Examples of arachidonic acid (A-D) and linoleic acid (E-H) derived lipid mediator average normalized signal intensities (NSI) acquired during 60 days for deep partial thickness burn wounds and full thickness burn wounds. Nb60 represents the values determined in the non-burned skin samples harvested at day 60. *Significant differences (p<0.05) between treatments in the corresponding time point.

## 4. Discussion and Conclusions

FSG contain high amounts of EPA and DHA but their direct involvement in the biochemical processes of wound healing has not been demonstrated. Based on the biological properties of EPA and DHA previously described in the literature [21, 22] we hypothesized that the two ω3-FA might affect the resolution of inflammation or other relevant wound healing stages through the formation of lipid mediators. In our study, we monitored the profiles of approximately 60 known lipid mediators with primarily pro-anti- and resolving inflammation properties. The treatment of burn wounds with FSG significantly affected both EPA and DHA metabolic pathways by increasing the amount of their mono-hydroxylated derivatives.

The analyzed biopsies were collected from DPT and FT wounds from a porcine burn model as outlined in **Figure 1**. Wound treatments followed standard procedures, normally applied in hospitals for similar types of wounds.[23] During this experiment, perfusion and closure rates and scar formation were also monitored and the results have been already reported elsewhere.[20] DPT showed quicker reepithelization in FSG treated wounds (statistically significant at day 14) with a faster subsidence of neutrophils (after 3 weeks), as opposed to fetal bovine dermis treated wounds that were still infiltrated with neutrophils at 4 weeks. Moreover, a persistent presence of macrophages was observed throughout the course of observation in fetal bovine dermis treated wounds while following FSG treatment, macrophage recruitment decreased after day 21 perhaps indicating a switch to a more progressive wound remodeling phase characterized by collagen reorganization and maturation. The data presented in this study are complementary to these observations and may serve to explain part of the mechanism that underlie the FSG mechanism of action.

Here, the changes to the profile of lipid content of the porcine biopsies were measured using UPLC-MS/MS (see supp. information) and their average NSI were calculated and compared at each time point. The UPLC-MS/MS method included compounds that belong to a wide spectrum of lipid mediators that belong to either EPA, DHA, AA, or LA metabolic pathway and can be considered as established biomarkers of the wound healing processes.[10, 24] Since wound healing is separated into four discrete yet overlapping phases [25] and the lipid mediators of interest were monitored for 60 days the concentration changes presented in **Figure 2** and the supplementary information span all these phases. The results showed that the majority of the measured lipid mediators had a concentration maximum two or three weeks (in day 14 or 21 respectively) after the creation of the wound independently of the wound type or treatment. These results conform with the general quantitative changes to lipid mediators during wound healing and reflect the multifunctional signaling properties of the lipid mediators during different overlapping wound healing phases.[5, 11] A more thorough temporal sampling profile is however required to capture the dynamics of lipid mediator profiles in each wound healing stage.

As opposed to other measured lipid mediators, most of the EPA and DHA derived mediators -principally the mono hydroxylated ones as well as the free EPA and DHA-were altered in the FSG treated wounds in the first 7 days of treatment in comparison to the other treatments used (**Figure 3 and 4**). These derivatives, although not always significantly different in both type of wounds, presented a trend of increased concentration in the FSG treated wounds that can be associated to the higher lipidic load of the graft. Based on our previous observations, [19] both EPA and DHA of the FSG lipid content are principally part of phosphatidylcholines and hence these alterations indicate the enzymatic release of ω3-FA by phospholipases, allowing secretion and further metabolism within the wound.

The increased presence of the mono-hydroxylated derivatives indicates activity of ALOX5, ALOX12, or ALOX15 and although the time course of the events may not be clear, these results support the importance of the ALOX enzymes in wound healing processes. The increased activity of the enzymes may be associated to the higher number of blood vessels observed in the FSG treated wounds during the first seven days of healing.[20] Further investigation of their activity may give more detailed insights into the way that external application of ω3-FA contributes to wound healing. In light of the trans cellular synthesis pathways required to generate these compounds [22] the results imply that immune cell recruitment into wounds treated with FSG is different than in non-FSG treated wounds. A more detailed immunological investigation focusing on white blood cells and cytokines would however be required to confirm this. Regarding other EPA and DHA hydroxylated derivatives measured in our study (e.g., 15-HEPE and 4-, 7-, 10-DHA), the observed changes of their amount in the FSG treated wounds corroborate the evidence of FSG derived EPA and DHA topical activity although our knowledge on their biological activity is limited.[26] Moreover, it is highly possible that more differences might be found in other EPA and DHA derived derivatives (e.g. ethanolamine or ceramide) if they were monitored.

Some special attention should be drawn to the increase of 18-HEPE and 17-HDHA after the first seven days in the wounds treated with FSG. Both molecules are precursors of the SPM superfamily which have been primarily associated with the resolution of inflammation and direct effects on epidermal keratinocytes,[27] promoting their migration.[28, 29] EPA and DHA are converted to 18-HEPE (acetylated COX2/CYP450 activity) and 17-HDHA (ALOX15 activity) respectively before they are eventually transformed to the corresponding resolvins through ALOX5 activity.[14] 18-HEPE and 17-HDHA are therefore biomarkers for the formation of the E-and D-series of resolvins as well as the activity of the implicated enzymes.[24] In the analyzed biopsies however, RvD1 as well as all the other measured SPMs (e.g., RvE1, RvE2, RvE3, RvD2, 7S-MaR1, PD1, and PDX) were not detected in any of the conditions. The absence of these compounds might be associated with the fact their formation is associated with the very early stages of the resolution of inflammation,[22] that are prior to the day that our first biopsy was collected. A much denser sampling during the first hours to days should be mandatory in order to exclude the formation of this compounds.

In addition to the observed differences in the amounts of EPA and DHA metabolites, significant changes were observed in the AA and LA pathways for the FTs. In this wound type, PGF2α and 13,14-dihydro-15-keto-PGF2α (AA pathway) and 13-HoDE and 13- HoTrE (LA pathway) were found to be significantly lower when treated with treatment 3 than with treatment 4. Among others, recent studies have shown that PGF2α and its keto-derivative (a stable metabolite of PGF2α that reflects in vivo PGF2α biosynthesis[30]) have been shown to play a significant role in pain modulation.[31] Similarly, both 13-HoDE and 13-HoTrE are part of a metabolic pathway that modulate the TprV1 receptor associated with pain.[32, 33] Although our results do not provide any conclusive information, they support a relationship between the FSG external application and pain modulation in FTs.

A limitation of our study consists that the biopsies were extracted as whole tissue and were not separated in layers. Thus, any information regarding the concentration of the compounds in the distinct layers was lost. This may explain why some significant differences between treatments were observed only for the FT, since deeper layers of the skin may recruit more macrophages and other cells that can readily metabolize the ω3-FA. Another limitation is that the cellular content in the wounds might be distinct due to the different biological material that each graft is made of. Thus, at some of the earlier time points when the punch biopsies were collected, part of the applied grafts could also be harvested, so that the observed chemical content could be affected by the different cellular ingrowth into the materials and in the case of cadaver skin, its content. This would be less relevant at late time points for fish skin in DPT as the graft breaks down over time and is integrated into the tissue. Moreover, a future study should be carried to determine the quantity of PLA2G2F, the secreted phospholipase with substrate preference for DHA-containing phospholipids, which is selectively expressed in the skin and regulates epidermal homeostasis.[34] Since both EPA and DHA are attached to the FSG in form of phosphatidylcholines, the presence of PLA2G2F may severely affect the amount of these ω3-FA that can enter the biochemical processes of wound healing. Finally, in this study the changes in the protein markers of inflammation (i.e. cytokines) were not assessed. This information, combined with the changes in cell ingrowth, protein biomarkers, and lipid biomarkers can a more mechanistic understanding of how FSG influences wound healing. For this, a more comprehensive -omics type study would be required.

In conclusion, our study demonstrated that the application of FSG to burn wounds results in higher formation of hydroxylated derivatives of both EPA (e.g., 18-HEPE) and DHA (e.g., 17-HDHA) when compared to other grafts of mammalian origin. The differences where observed in the first seven days after the application of the treatment indicating the interaction of the graft during the early stage of the healing process. More research is required to elucidate the exact mechanisms through which the ω3-FA found on FSG take part in these changes. The findings however clearly demonstrate that FSG influences lipid mediator biosynthesis in burn wound healing and might explain part of the beneficiary properties of FSG towards wound healing.

## 5. Acknowledgments

The authors would like to thank Dr. Hilmar Kjartansson M.D. and Ragnhildur Þórarinsdóttir for their contribution to interpretation of the data and help with figures. The views expressed in this article are those of the authors and do not reflect the official policy or position of the U.S. Army Medical Department, Department of the Army, DoD, or the U.S. Government.

## Abbreviations

AA: arachidonic acid
wwDHA: docosahexaenoic acid
DPT: deep partial thickness burn wounds
EPA: eicosapentaenoic acid
FSG: acellular cod fish skin grafts
FT: full thickness burn wounds
HETE: hydroxy- eicosatetraenoic acids
KETE: keto- eicosatetraenoic acids
LA: linoleic acid
MeOH: methanol
NSI: normalized standard intensity
RvD1: Resolvin D1
SPMs: specialized pro-resolving lipid mediators
ω3-FA: omega-3 fatty acids

## Author contributions

**Aristotelis Kotronoulas:** Conceptualization, Methodology, Formal analysis, Investigation, Data Curation, Writing -Original Draft, Project administration **Sigurður Trausti Karvelsson:** Investigation, Writing -Review & Editing **Marieke Heijink:** Investigation, Writing -Review & Editing **Randolph Stone II:** Investigation, Writing -Review & Editing **Martin Giera:** Resources, Writing -Review & Editing, Supervision **Ottar Rolfsson:** Conceptualization, Methodology, Resources, Writing -Review & Editing, Supervision, Funding acquisition

## References

[1] V. Puri, N.A. Khare, M.V. Chandramouli, N. Shende, S. Bharadwaj, Comparative Analysis of Early Excision and Grafting vs Delayed Grafting in Burn Patients in a Developing Country, J Burn Care Res, 37 (2016) 278–282.

[2] V.C. van der Veen, M.B. van der Wal, M.C. van Leeuwen, M.M. Ulrich, E. Middelkoop, Biological background of dermal substitutes, Burns, 36 (2010) 305–321.

[3] T. Yamamoto, H. Iwase, T.W. King, H. Hara, D.K.C. Cooper, Skin xenotransplantation: Historical review and clinical potential, Burns, 44 (2018) 1738–1749.

[4] C.K. Yang, T.O. Polanco, J.C. Lantis, 2nd, A Prospective, Postmarket, Compassionate Clinical Evaluation of a Novel Acellular Fish-skin Graft Which Contains Omega-3 Fatty Acids for the Closure of Hard-to-heal Lower Extremity Chronic Ulcers, Wounds, 28 (2016) 112–118.

[5] K. Alam, S.L.A. Jeffery, Acellular Fish Skin Grafts for Management of Split Thickness Donor Sites and Partial Thickness Burns: A Case Series, Mil Med, 184 (2019) 16–20.

[6] B. Dorweiler, T.T. Trinh, F. Dunschede, C.F. Vahl, E.S. Debus, M. Storck, H. Diener, The marine Omega3 wound matrix for treatment of complicated wounds: A multicenter experience report, Gefasschirurgie, 23 (2018) 46–55.

[7] B.T. Baldursson, H. Kjartansson, F. Konradsdottir, P. Gudnason, G.F. Sigurjonsson, S.H. Lund, Healing rate and autoimmune safety of full-thickness wounds treated with fish skin acellular dermal matrix versus porcine small-intestine submucosa: a noninferiority study, Int J Low Extrem Wounds, 14 (2015) 37–43.

[8] S. Magnusson, B.T. Baldursson, H. Kjartansson, O. Rolfsson, G.F. Sigurjonsson, Regenerative and Antibacterial Properties of Acellular Fish Skin Grafts and Human Amnion/Chorion Membrane: Implications for Tissue Preservation in Combat Casualty Care, Mil Med, 182 (2017) 383–388.

[9] R. Zeng, C. Lin, Z. Lin, H. Chen, W. Lu, C. Lin, H. Li, Approaches to cutaneous wound healing: basics and future directions, Cell Tissue Res, 374 (2018) 217–232.

[10] N.X. Landen, D. Li, M. Stahle, Transition from inflammation to proliferation: a critical step during wound healing, Cell Mol Life Sci, 73 (2016) 3861–3885.

[11] J.R. Silva, B. Burger, C.M.C. Kuhl, T. Candreva, M.B.P. Dos Anjos, H.G. Rodrigues, Wound Healing and Omega-6 Fatty Acids: From Inflammation to Repair, Mediators Inflamm, 2018 (2018) 2503950.

[12] M. Spite, J. Claria, C.N. Serhan, Resolvins, specialized proresolving lipid mediators, and their potential roles in metabolic diseases, Cell Metab, 19 (2014) 21–36.

[13] R. Menon, P. Krzyszczyk, F. Berthiaume, Pro-Resolution Potency of Resolvins D1, D2 and E1 on Neutrophil Migration and in Dermal Wound Healing, Nano Life, 7 (2017).

[14] C.N. Serhan, Pro-resolving lipid mediators are leads for resolution physiology, Nature, 510 (2014) 92–101.

[15] O. Kuda, Bioactive metabolites of docosahexaenoic acid, Biochimie, 136 (2017) 12–20.

[16] E.L. Campbell, N.A. Louis, S.E. Tomassetti, G.O. Canny, M. Arita, C.N. Serhan, S.P. Colgan, Resolvin E1 promotes mucosal surface clearance of neutrophils: a new paradigm for inflammatory resolution, FASEB J, 21 (2007) 3162–3170.

[17] P.C. Calder, Omega-3 fatty acids and inflammatory processes, Nutrients, 2 (2010) 355–374.

[18] J.C. McDaniel, K. Massey, A. Nicolaou, Fish oil supplementation alters levels of lipid mediators of inflammation in microenvironment of acute human wounds, Wound Repair Regen, 19 (2011) 189–200.

[19] A. Kotronoulas, H.S. Jonasdottir, R.S. Sigurethardottir, S. Halldorsson, G.G. Haraldsson, O. Rolfsson, Wound healing grafts: Omega-3 fatty acid lipid content differentiates the lipid profiles of acellular Atlantic cod skin from traditional dermal substitutes, J Tissue Eng Regen Med, 14 (2020) 441–451.

[20] R. Stone, 2nd, E.C. Saathoff, D.A. Larson, J.T. Wall, N.A. Wienandt, S. Magnusson, H. Kjartansson, S. Natesan, R.J. Christy, Accelerated Wound Closure of Deep Partial Thickness Burns with Acellular Fish Skin Graft, Int J Mol Sci, 22 (2021).

[21] P.C. Calder, Marine omega-3 fatty acids and inflammatory processes: Effects, mechanisms and clinical relevance, Biochim Biophys Acta, 1851 (2015) 469–484.

[22] C.N. Serhan, S.K. Gupta, M. Perretti, C. Godson, E. Brennan, Y. Li, O. Soehnlein, T. Shimizu, O. Werz, V. Chiurchiu, A. Azzi, M. Dubourdeau, S.S. Gupta, P. Schopohl, M. Hoch, D. Gjorgevikj, F.M. Khan, D. Brauer, A. Tripathi, K. Cesnulevicius, D. Lescheid, M. Schultz, E. Sarndahl, D. Repsilber, R. Kruse, A. Sala, J.Z. Haeggstrom, B.D. Levy, J.G. Filep, O. Wolkenhauer, The Atlas of Inflammation Resolution (AIR), Mol Aspects Med, (2020) 100893.

[23] T.L. Khoo, A.S. Halim, A.Z. Saad, A.A. Dorai, The application of glycerol-preserved skin allograft in the treatment of burn injuries: an analysis based on indications, Burns, 36 (2010) 897–904.

[24] C.N. Serhan, N. Chiang, J. Dalli, New pro-resolving n-3 mediators bridge resolution of infectious inflammation to tissue regeneration, Mol Aspects Med, 64 (2018) 1–17.

[25] J.M. Reinke, H. Sorg, Wound repair and regeneration, Eur Surg Res, 49 (2012) 35–43.

[26] R.G. Snodgrass, B. Brune, Regulation and Functions of 15-Lipoxygenases in Human Macrophages, Front Pharmacol, 10 (2019) 719.

[27] J. Hellmann, B.E. Sansbury, B. Wong, X. Li, M. Singh, K. Nuutila, N. Chiang, E. Eriksson, C.N. Serhan, M. Spite, Biosynthesis of D-Series Resolvins in Skin Provides Insights into their Role in Tissue Repair, J Invest Dermatol, 138 (2018) 2051–2060.

[28] M.L. Usui, J.N. Mansbridge, W.G. Carter, M. Fujita, J.E. Olerud, Keratinocyte migration, proliferation, and differentiation in chronic ulcers from patients with diabetes and normal wounds, J Histochem Cytochem, 56 (2008) 687–696.

[29] M.A. Seeger, A.S. Paller, The Roles of Growth Factors in Keratinocyte Migration, Adv Wound Care (New Rochelle), 4 (2015) 213–224.

[30] E. Ricciotti, G.A. FitzGerald, Prostaglandins and inflammation, Arterioscler Thromb Vasc Biol, 31 (2011) 986–1000.

[31] S. Kunori, S. Matsumura, T. Mabuchi, S. Tatsumi, Y. Sugimoto, T. Minami, S. Ito, Involvement of prostaglandin F 2 alpha receptor in ATP-induced mechanical allodynia, Neuroscience, 163 (2009) 362–371.

[32] M. Alsalem, A. Wong, P. Millns, P.H. Arya, M.S. Chan, A. Bennett, D.A. Barrett, V. Chapman, D.A. Kendall, The contribution of the endogenous TRPV1 ligands 9-HODE and 13-HODE to nociceptive processing and their role in peripheral inflammatory pain mechanisms, Br J Pharmacol, 168 (2013) 1961–1974.

[33] V. Vangaveti, V. Shashidhar, F. Collier, J. Hodge, C. Rush, U. Malabu, B. Baune, R.L. Kennedy, 9-and 13-HODE regulate fatty acid binding protein-4 in human macrophages, but does not involve HODE/GPR132 axis in PPAR-gamma regulation of FABP4, Ther Adv Endocrinol Metab, 9 (2018) 137–150.

[34] K. Yamamoto, Y. Miki, M. Sato, Y. Taketomi, Y. Nishito, C. Taya, K. Muramatsu, K. Ikeda, H. Nakanishi, R. Taguchi, N. Kambe, K. Kabashima, G. Lambeau, M.H. Gelb, M. Murakami, The role of group IIF-secreted phospholipase A2 in epidermal homeostasis and hyperplasia, J Exp Med, 212 (2015) 1901–1919.

