## Supplementary information for "Lipid mediator profile of burn wound healing: Acellular cod fish skin grafts promote the formation of EPA and DHA derived lipid mediators during the first seven days"

**Table S1:** Summary of the MS/MS detection parameters for all lipid mediators. *(Q1 Mass):* parent mass used for the first quadrupole, (*Q3 Mass):* fragment mass used for the third quadrupole, *(RT):* retention time, *(DP):* declustering potential, *(EP):* entrance potential, *(CE):* collision energy and *(CXP):* collision cell exit potential

| ID | Q1 Mass | Q3 Mass | RT (min) | DP (volts) | EP (volts) | CE (volts) | CXP (volts) |
|---|---|---|---|---|---|---|---|
| 10-HDHA | 343.1 | 153 | 7.9 | -25 | -10 | -20 | -15 |
| 11(12)EET | 318.9 | 166.9 | 8.2 | -90 | -10 | -18 | -19 |
| 11-HETE | 319.1 | 167 | 7.9 | -70 | -10 | -22 | -15 |
| 12-HEPE | 317 | 179 | 7.6 | -60 | -10 | -18 | -17 |
| 12-HETE | 319.1 | 179 | 7.9 | -65 | -10 | -20 | -23 |
| 12-KETE | 317 | 153 | 7.85 | -60 | -10 | -22 | -9 |
| 13,14dihydro-15keto-PGF2a | 353.1 | 195 | 5.4 | -110 | -10 | -32 | -11 |
| 13-HoDE | 295 | 194.9 | 7.7 | -110 | -10 | -24 | -21 |
| 13-HoTrE | 293 | 195 | 7.4 | -45 | -10 | -24 | -19 |
| 14(15)EET | 319 | 218.9 | 8.05 | -5 | -10 | -16 | -55 |
| 14(S)-HDHA | 343.1 | 204.9 | 7.95 | -60 | -10 | -18 | -27 |
| 14,15-diHETE | 335.1 | 207 | 6.95 | -65 | -10 | -24 | -21 |
| 15Deoxy PGJ2 | 315 | 203 | 7.3 | -50 | -10 | -28 | -19 |
| 15-HEPE | 317.1 | 219 | 7.5 | -65 | -10 | -18 | -19 |
| 15-HETE | 319.1 | 219.1 | 7.8 | -55 | -10 | -18 | -9 |
| 15-HETE d8 | 327.2 | 226 | 7.8 | -85 | -10 | -18 | -11 |
| 15-KETE | 317 | 113 | 7.75 | -10 | -10 | -22 | -5 |
| 15-Keto-PGE2 | 349 | 234.9 | 4.45 | -65 | -10 | -20 | -13 |
| 17F2 Dihomo IsoP | 381.1 | 318.9 | 6.05 | -115 | -10 | -32 | -41 |
| 17-HDHA | 343.1 | 245 | 7.85 | -65 | -10 | -16 | -15 |
| 17-OH-DH-HETE | 347.1 | 247 | 8.2 | -110 | -10 | -22 | -27 |
| 18-HEPE | 317.1 | 259 | 7.4 | -5 | -10 | -16 | -7 |
| 18R-RvE3 | 333.1 | 245 | 7.1 | -55 | -10 | -18 | -23 |
| 18S-RvE3 | 333.1 | 245.2 | 6.7 | -25 | -10 | -16 | -17 |
| 19(20)-EpDPA | 343.1 | 281.1 | 8.05 | -70 | -10 | -16 | -11 |
| 19,20-DiHDPA | 361.1 | 273 | 7.35 | -55 | -10 | -22 | -15 |
| 20-HETE_2 | 319 | 289.1 | 7.67 | -70 | -10 | -24 | -15 |
| 20-OH LTB4 | 351.1 | 195 | 4 | -60 | -10 | -24 | -17 |
| 4-HDHA | 343.1 | 101 | 8.15 | -50 | -10 | -18 | -9 |
| 5,15-diHETE | 335 | 173.1 | 6.75 | -55 | -10 | -20 | -11 |
| 5-HEPE | 317 | 114.9 | 7.7 | -55 | -10 | -18 | -11 |
| 5-HETE | 319.1 | 115 | 8 | -65 | -10 | -18 | -11 |
| 5-KETE | 317 | 203.1 | 8.05 | -70 | -10 | -24 | -11 |
| 6t,12epi-LTB4 | 335.1 | 194.9 | 6.8 | -80 | -10 | -22 | -25 |
| 6-trans-LTB4 | 335.1 | 194.9 | 6.65 | -105 | -10 | -22 | -11 |
| 7,17-DiHDPA | 361.1 | 198.9 | 7 | -45 | -10 | -26 | -23 |
| 7-HDHA | 343.1 | 141.1 | 7.95 | -85 | -10 | -18 | -23 |
| 7S-MaR1 | 359.1 | 249.9 | 6.6 | -20 | -10 | -20 | -19 |

### Table S1 (continuation)

| ID | Q1 Mass | Q3 Mass | RT (min) | DP (volts) | EP (volts) | CE (volts) | CXP (volts) |
|---|---|---|---|---|---|---|---|
| 8(9)EET | 319 | 154.9 | 8.2 | -60 | -10 | -18 | -13 |
| 8-HETE | 319.1 | 154.9 | 7.9 | -70 | -10 | -20 | -19 |
| 8-iso-PGE2 | 351.1 | 271 | 4.6 | -5 | -10 | -24 | -19 |
| 8-iso-PGF2a | 353.1 | 193 | 4.45 | -135 | -10 | -34 | -11 |
| 8-iso-PGF2alpha-d4 | 357.3 | 197 | 4.45 | -110 | -10 | -34 | -20 |
| 8S,15S-diHETE | 335.1 | 207.9 | 6.7 | -55 | -10 | -22 | -17 |
| 9-HoDE | 295 | 171 | 7.7 | -130 | -10 | -22 | -7 |
| 9-HoTrE | 293 | 170.9 | 7.35 | -75 | -10 | -20 | -15 |
| AA | 303 | 205.1 | 8.75 | -155 | -10 | -20 | -11 |
| AdA | 331.1 | 233 | 9.05 | -130 | -10 | -22 | -11 |
| ALA/GLA | 277 | 233 | 8.55 | -90 | -10 | -22 | -29 |
| AT-LXA4 | 351.1 | 114.9 | 5.6 | -20 | -10 | -22 | -11 |
| AT-RvD1 | 375 | 215 | 5.7 | -50 | -10 | -26 | -11 |
| DGLA | 305.1 | 261.2 | 8.95 | -85 | -10 | -22 | -13 |
| DHA | 327.1 | 229.2 | 8.75 | -115 | -10 | -18 | -11 |
| DHAd5 | 332 | 288.1 | 8.75 | -75 | -10 | -16 | -13 |
| DPAn-3 | 329.1 | 231.1 | 8.95 | -50 | -10 | -20 | -17 |
| EPA | 301 | 202.9 | 8.55 | -125 | -10 | -18 | -21 |
| LA | 279 | 261 | 8.8 | -115 | -10 | -28 | -13 |
| Leukotriene B4 | 335.1 | 195 | 6.9 | -65 | -10 | -22 | -21 |
| Leukotriene B4 d4 | 339.1 | 196.9 | 6.9 | -70 | -10 | -22 | -19 |
| LTD4 | 495.1 | 177 | 6.65 | -70 | -10 | -28 | -19 |
| LTE4 | 438.1 | 333.1 | 6.95 | -55 | -10 | -26 | -15 |
| LXA4 | 351.1 | 114.8 | 5.45 | -40 | -10 | -20 | -11 |
| LXB4 | 351.1 | 220.9 | 5.05 | -60 | -10 | -22 | -13 |
| MaR1_2 | 359.2 | 250.2 | 6.95 | -65 | -10 | -20 | -13 |
| PD1 | 359.1 | 153 | 6.9 | -70 | -10 | -22 | -9 |
| PDX | 359.1 | 153 | 6.8 | -70 | -10 | -22 | -9 |
| PGD2 | 351.1 | 233 | 4.95 | -30 | -10 | -16 | -13 |
| PGE2_2 | 351.2 | 271.1 | 4.85 | -50 | -10 | -22 | -21 |
| PGE2-d4 | 355.1 | 193 | 4.85 | -50 | -10 | -26 | -17 |
| PGF2a | 353.1 | 193 | 5.15 | -80 | -10 | -34 | -11 |
| PGJ2 | 333 | 271 | 6.05 | -30 | -10 | -22 | -17 |
| RvD1 | 375.1 | 215 | 5.55 | -50 | -10 | -26 | -11 |
| RvD2 | 375.1 | 277.1 | 5.25 | -60 | -10 | -18 | -15 |
| RvE1 | 349.1 | 195 | 3.8 | -95 | -10 | -22 | -13 |
| RvE2 | 333.1 | 114.9 | 6.1 | -35 | -10 | -18 | -15 |
| TxB2 | 369.1 | 169 | 4.6 | -55 | -10 | -24 | -15 |

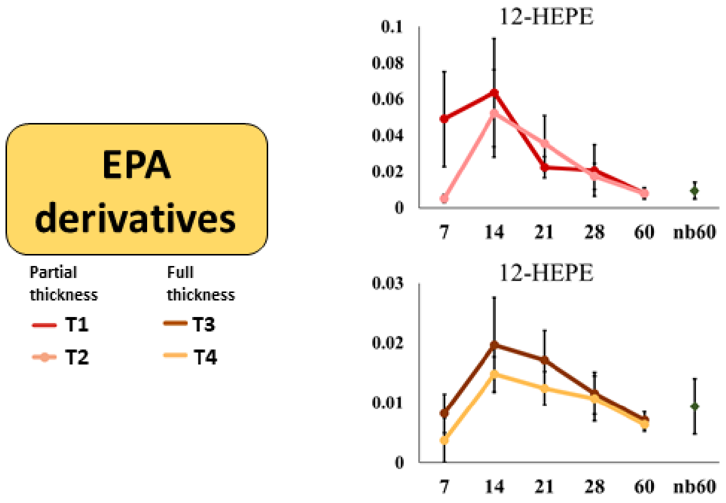


**Figure S1:** EPA derived lipid mediator profiles acquired overs 60 days of wound healing for PTBW and FTBW. Intensities of lipids are displayed as average normalized signal intensities (NSI).

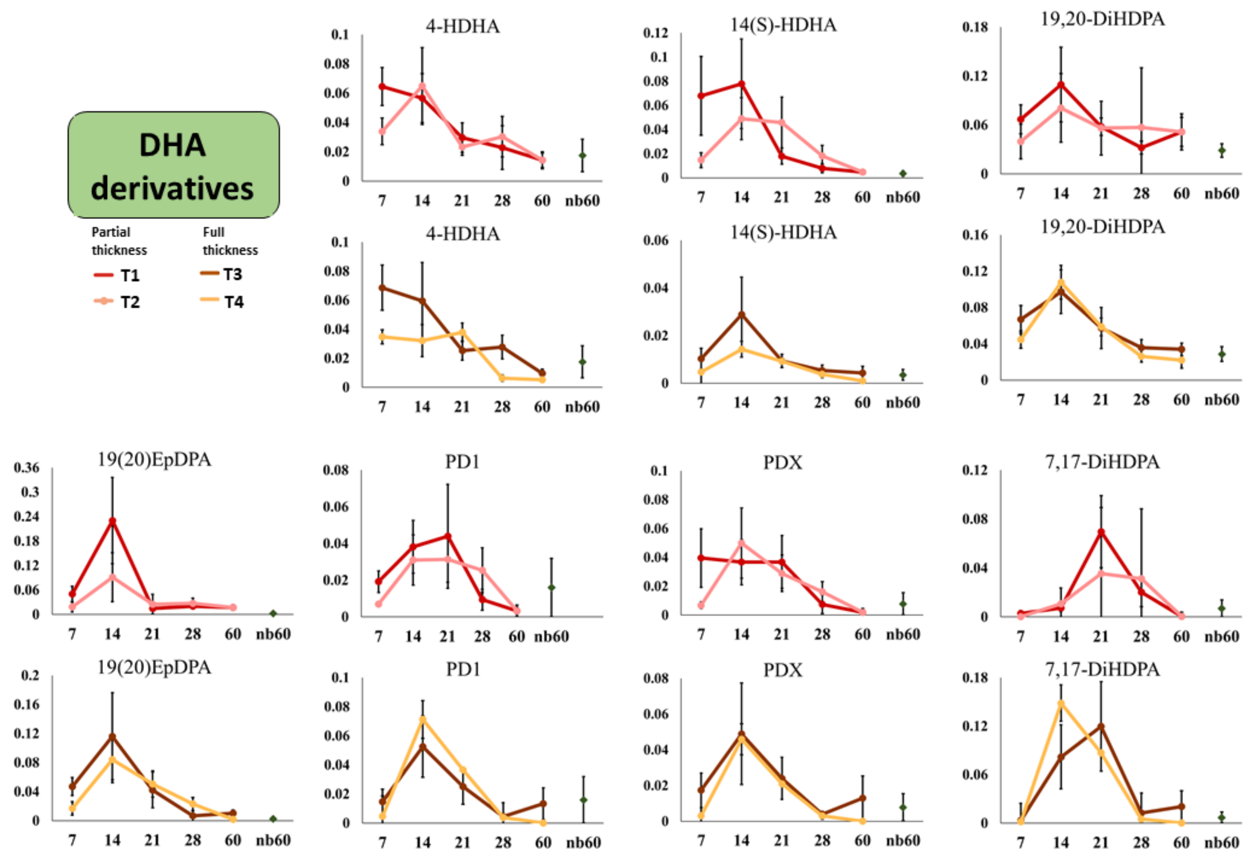


**Figure S2:** DHA derived lipid mediator profiles acquired overs 60 days of wound healing for PTBW and FTBW. Intensities of lipids are displayed as average normalized signal intensities (NSI).

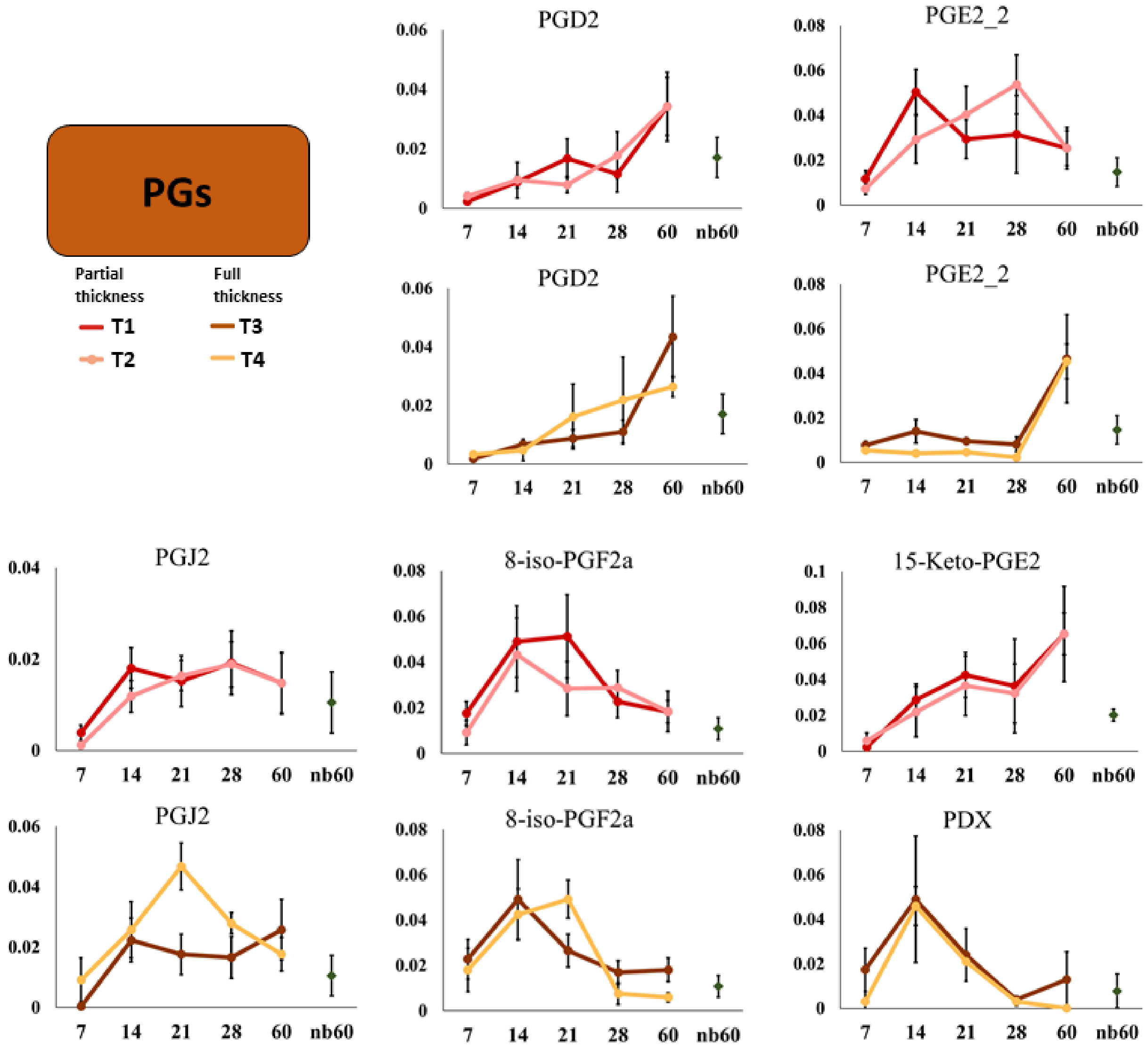


**Figure S3:** Prostaglandin (PGs) profiles acquired overs 60 days of wound healing for PTBW and FTBW. Intensities of lipids are displayed as average normalized signal intensities (NSI).

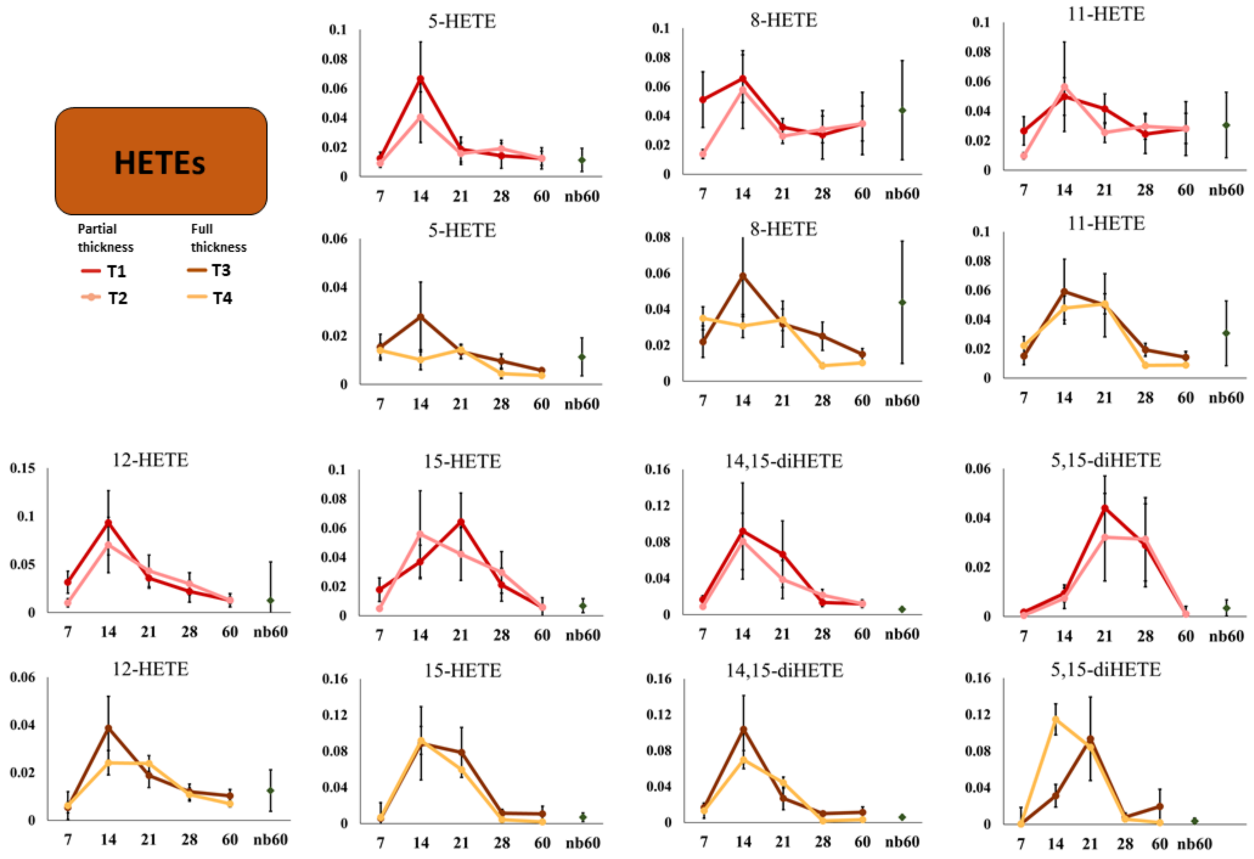


**Figure S4:** Hydroxyeicosatetraenoic acids (HETEs) profiles acquired overs 60 days of wound healing for PTBW and FTBW. Intensities of lipids are displayed as average normalized signal intensities (NSI).

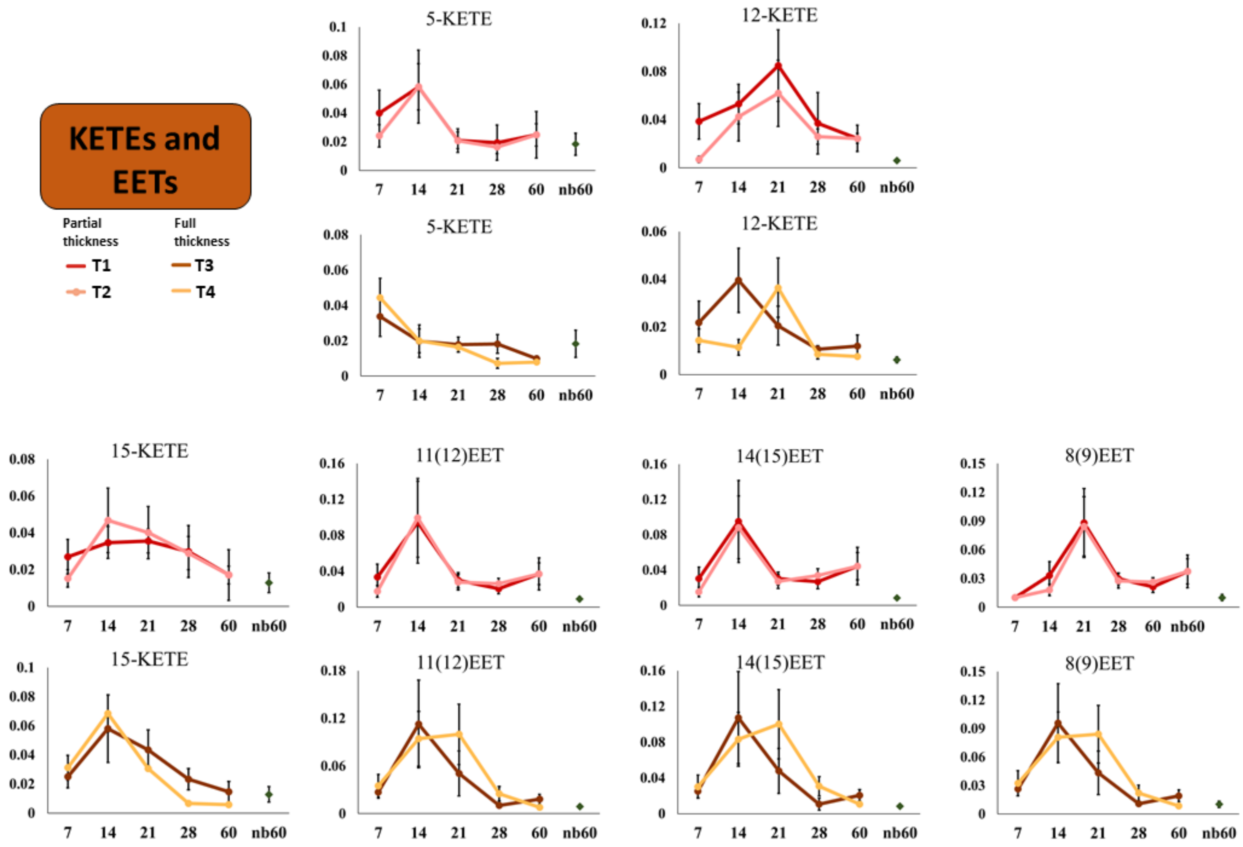


**Figure S5:** Oxoeicosatetraenoic (KETEs) and epoxyeicosatrienoic acids (EETs) profiles acquired overs 60 days of wound healing for PTBW and FTBW. Intensities of lipids are displayed as average normalized signal intensities (NSI).

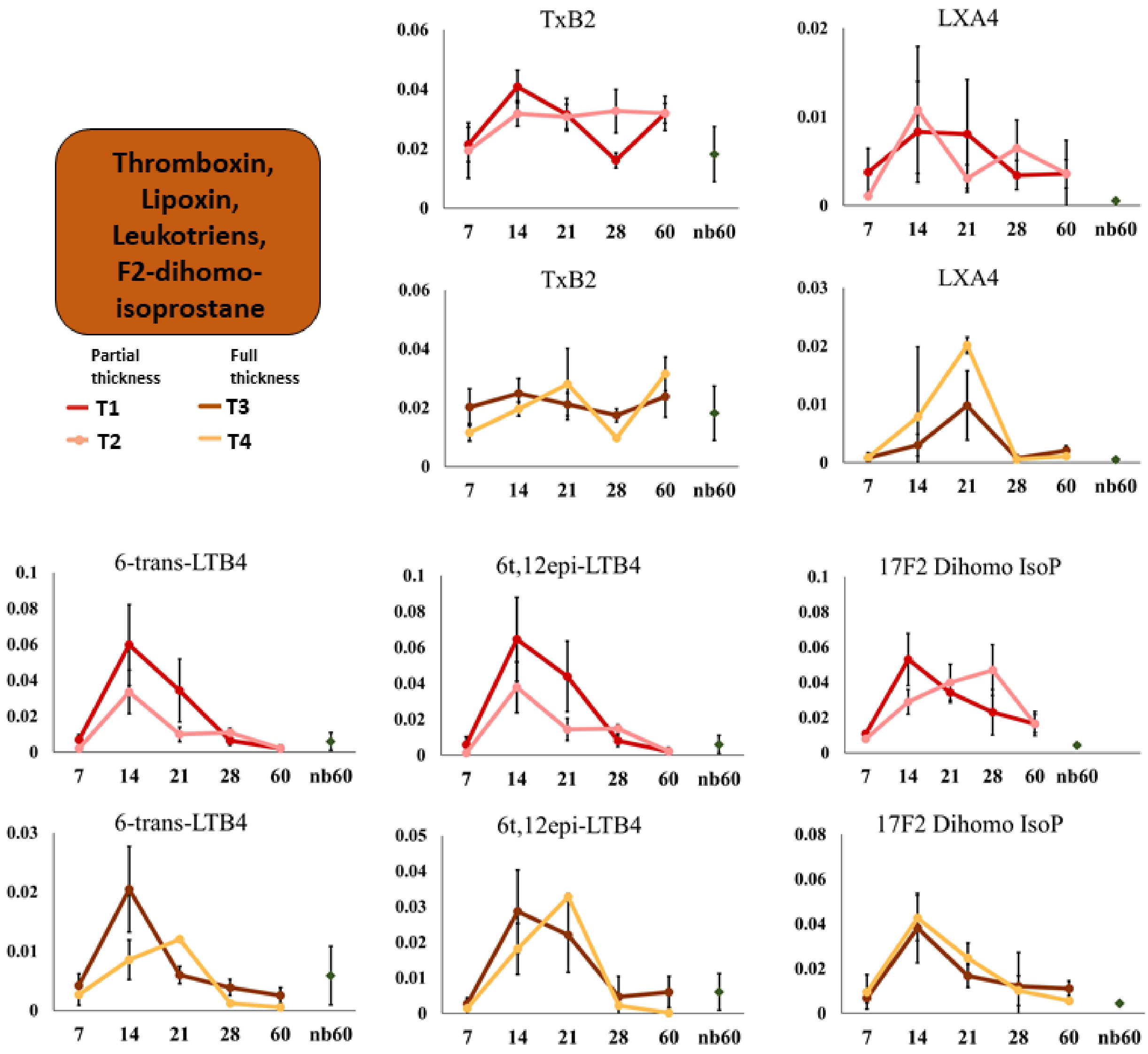


**Figure S6:** Thromboxine B4, leukotrienes and F2-dihomo-isoprostane profiles acquired overs 60 days of wound healing for PTBW and FTBW. Intensities of lipids are displayed as average normalized signal intensities (NSI).

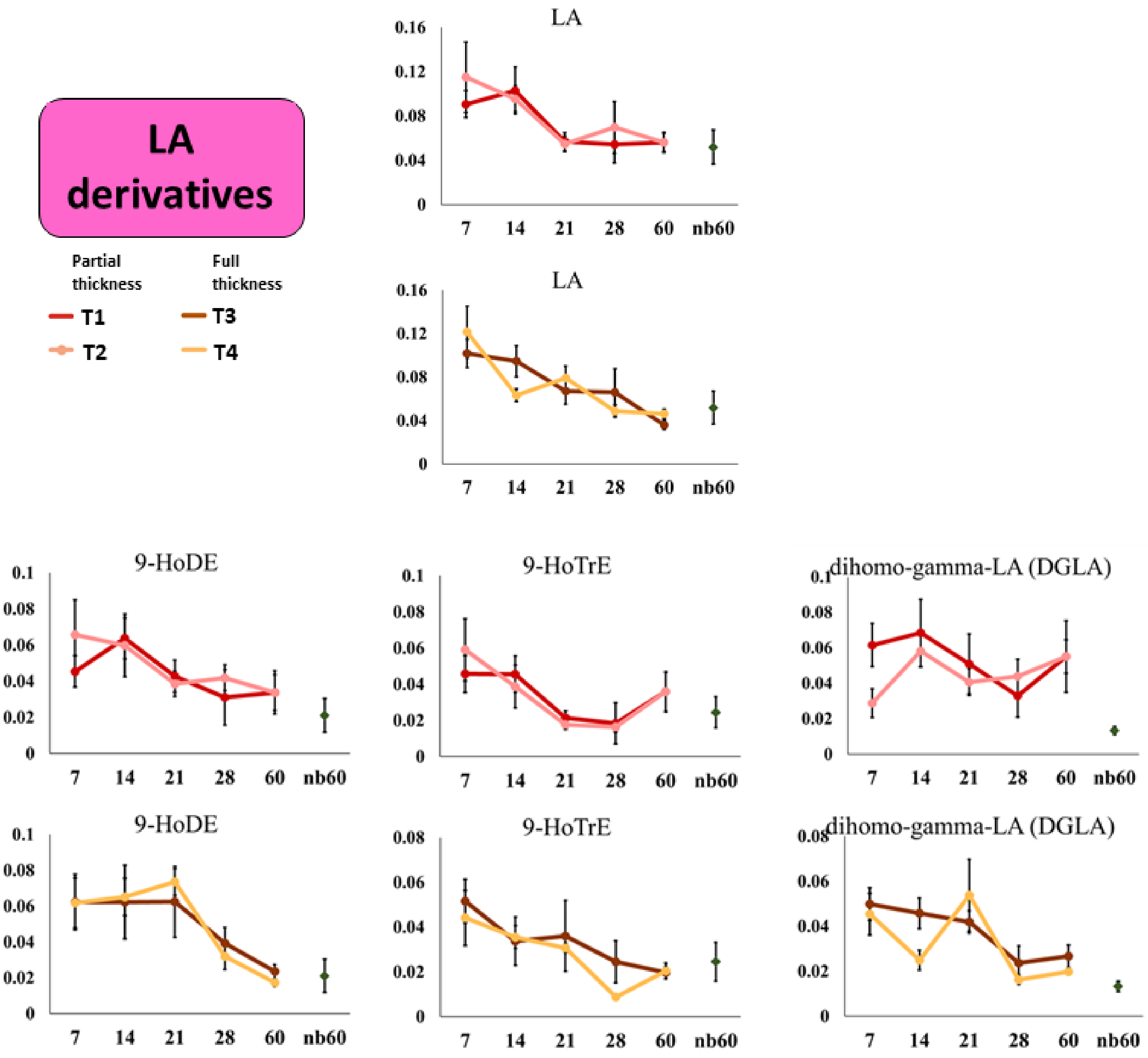


**Figure S7:** Linoleic acid (LA) derived lipid mediator profiles acquired overs 60 days of wound healing for PTBW and FTBW. Intensities of lipids are displayed as average normalized signal intensities (NSI).

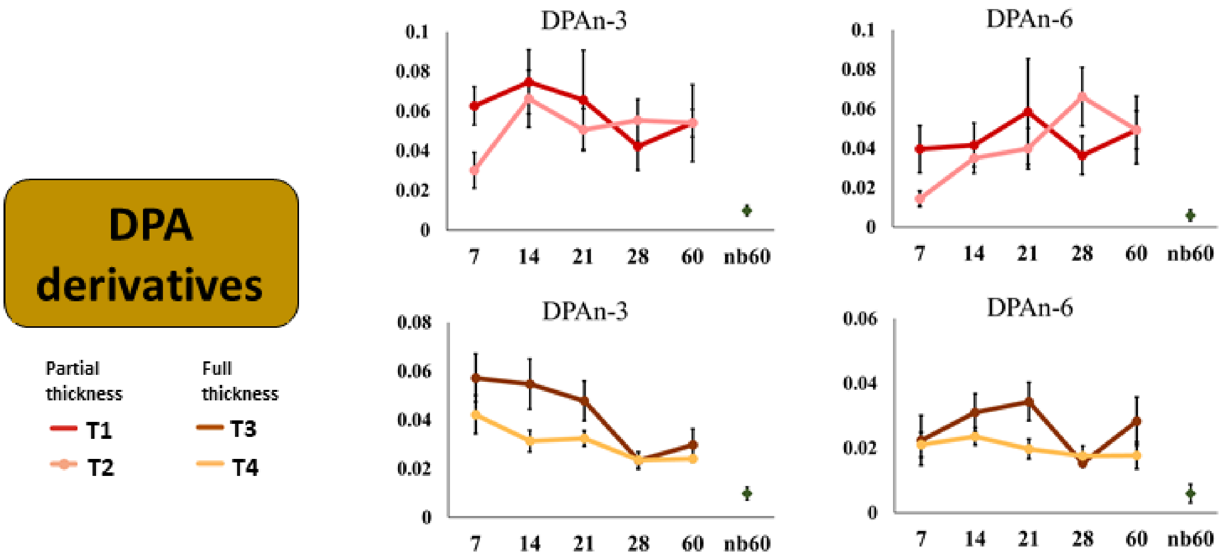


**Figure S8:** Docosapentaenoic acid (DPA) derived lipid mediator profiles acquired overs 60 days of wound healing for PTBW and FTBW. Intensities of lipids are displayed as average normalized signal intensities (NSI).
